# Understanding the dynamics of scaffold-mediated signaling

**DOI:** 10.1101/167205

**Authors:** Ryan Suderman, Addison Schauer, Eric J. Deeds

**Affiliations:** Center for Computational Biology, University of Kansas, Lawrence KS 66045, USA; Department of Molecular Biosciences, University of Kansas, Lawrence KS 66045, USA

**Keywords:** scaffold proteins, kinase cascades, signaling dynamics, crosstalk, rule-based modeling

## Abstract

Many signaling networks involve scaffold proteins that bind multiple kinases in kinase cascades. While scaffolds play a fundamental role in regulating signaling, few hypotheses regarding their function have been rigorously examined. Here, we used dynamical models of scaffold signaling to investigate the impact scaffolds have on network behavior. We considered two paradigms of scaffold assembly: as either the nucleation point for assembly of discrete multi-subunit proteins (the machine paradigm) or a platform upon which kinases independently associate (the ensemble paradigm). We found that several well-accepted hypotheses regarding the role of scaffolds in regulating signal response either do not hold or depend critically on the assembly paradigm employed. In addition to providing novel insights into the function of scaffold proteins, our work suggests experiments that could distinguish between assembly paradigms. Our findings should also inform attempts to target scaffold proteins for therapeutic intervention and the design of scaffolds for synthetic biology.

## Introduction

Intracellular signaling networks form the basis for cellular adaptation to the environment, and kinase cascades are a common motif in these networks, particularly in eukaryotes (1, 2). Interestingly, many of these cascades involve a dedicated “scaffold protein,” which often have no catalytic activity themselves, but rather serve as a multivalent nucleation point for the assembly of signaling complexes (1, 3, 4). While scaffolds are common, there are clear examples of kinase cascades that function without them (1); this has led to a wide array of hypotheses regarding the functional role scaffolds play in the cascades in which they are found (3, 4). For instance, many have argued that scaffolds prevent signal amplification, based on the intuition that stoichiometric limitations imposed by the scaffold should limit activation of downstream species (3, 4). Others have speculated that scaffolds serve to prevent unwanted *crosstalk* between pathways, by sequestering kinases that are shared by two cascades onto a physical platform specific to one of them (5, 6). Despite the fact that scaffold proteins have been the subject of numerous theoretical and experimental studies (7-10), surprisingly few of these hypotheses have ever been explored in a rigorous way. Nonetheless, many of these ideas (particularly the concept that scaffolds limit or prevent signal amplification) have become widely accepted within the field (3, 4).

There are, of course, exceptions to the above statement, and one of the most prominent of these is *combinatorial inhibition*, a phenomenon similar to the *prozone effect* observed in immune response, in which excess scaffold concentration inhibits response to signal (9). The capacity for scaffold proteins to induce this effect was first explored computationally by Levchenko and co-workers (9) and was later confirmed experimentally in the yeast pheromone signaling network, which involves one of the most well-characterized MAP kinase cascades organized on the scaffold protein Ste5 (11, 12). Another computational study indicated that scaffolds are capable of preventing signal attenuation in kinase cascades that exhibit strong phosphatase activity (8). In particular, Locasale *et al.* found that increasing the binding affinity between kinases and the scaffold correspondingly increases the probability of activation events, since kinases are more likely to be located near one another on a scaffold. While this could represent one function of scaffold proteins, the authors modeled substrate activation/deactivation using instantaneous collision events, and so it is unclear how phenomena such as direct substrate binding by kinases, or enzyme saturation, might influence their results (8). Optimal binding to the scaffold and the effect of scaffold-enzyme association on the rate of catalysis have also been explored computationally in the context of scaffold-based cascades (13). Regardless, many prevailing hypotheses regarding scaffold function have yet to be investigated in detail (3).

One barrier to developing a general understanding of scaffold function is the fact that it is currently unclear exactly how kinases assemble onto the scaffold. Most representations of scaffold-based cascades in the literature summarize the relevant interactions by drawing all the kinases simultaneously bound to the scaffold (14). This is evocative of the orderly assembly of a machine-like “signalosome” with a well-defined composition and quaternary structure. Existing computational models of scaffold assembly, however, usually assume that binding to the scaffold is independent; in other words, the binding of one kinase to the scaffold does not influence the binding probability of other kinases (15, 16). One consequence of independent binding, however, is *combinatorial complexity*: as the number of binding partners of the scaffold grows (call this number “*N*”), the number of possible distinct molecular species increases as 2^*N*^. We recently showed that, given scaffold dimerization and the many phosphorylation states of the kinases themselves, the interactions involving the Ste5 scaffold in the yeast pheromone network can generate over 3 billion distinct biochemical species (14). If binding is (largely) independent, we found that the common representation of a fully assembled scaffold complex actually never forms during simulations of signaling dynamics; instead, signaling tends to proceed via a heterogeneous *ensemble* of protein complexes (14, 17). Formation of a more machine-like structure with all of the relevant proteins simultaneously bound to the scaffold requires specific hierarchical assembly constraints (14).

There is currently little experimental evidence regarding whether any given scaffold protein nucleates the formation of ensembles or machines *in vivo*. Interestingly, in our simulations of the pheromone network we found that machines and ensembles can exhibit very different behaviors; for instance, the classical result of combinatorial inhibition is only possible in ensemble-like signaling, at least in the Ste5 cascade (12, 14). This is similar to the reduction in combinatorial inhibition that had been observed with increasing cooperativity between the effectors that bind the scaffold (13). In this work, we constructed a series of computational models in order to systematically understand how scaffold-based cascades differ from cascades where there is no scaffold, and how ensemble-like signaling differs from signaling through machine-like structures. Our investigation of these models revealed that a number of seemingly intuitive and well-accepted ideas about scaffold function *do not necessarily hold*. For instance, while ensembles tend to have slightly less amplification than cascades without a scaffold, they can still amplify signals by over 100-fold, depending on the strength of the input signal. Machine-like assembly results in amplification equivalent to, or even greater than, that observed in solution cascades, implying that the existence of a scaffold within a cascade is by no means a guarantee that signal amplification will not occur. Scaffolds also do not necessarily prevent crosstalk: in ensemble models, we found that crosstalk is reduced, but not eliminated, when two cascades share a kinase but have distinct scaffolds. While machine-like scaffolds can prevent one cascade from inadvertently activating another, we found that activation of one pathway can actually decrease the activity of another in this model, indicating a potential for crosstalk even in that case.

These results underscore one of our key findings: in many cases, the assembly mechanism employed by the scaffold matters more than the presence of the scaffold itself. This implies that characterizing the assembly pathway is necessary for understanding the functional role of a scaffold within a signaling network. Since experimental work has generally not explored this aspect of scaffold dynamics, this is clearly an important area for future investigation. Our results also imply that many scaffold functions are mutually exclusive between paradigms, which could help to constrain hypotheses on scaffold function. As mentioned above, overexpressing the scaffold Ste5 in yeast results in significant combinatorial inhibition, which implies at least some ensemble-like character in Ste5 signaling (12). One component of the Ste5 cascade is the MAPKK Ste11, which is shared between the Ste5-based pheromone cascade and the High Osmolarity Glycerol (HOG) pathway in yeast, which is based on a different scaffold Pbs2. One of the main hypotheses regarding Ste5 function is the prevention of crosstalk between these two pathways (5); since ensemble-like scaffolds cannot prevent crosstalk, however, that is unlikely to be Ste5’s role in the network.

While more work is clearly necessary to fully characterize specific scaffolds’ assembly pathways and functions, the above highlights how the systematic modeling approach taken in this work can inform our interpretation of experimental data. In addition to suggesting experiments that could help constrain scaffold assembly mechanisms and function, our findings also have important implications for synthetic biology (3, 7) and the development of cancer therapies involving dysregulation of signaling cascades involving scaffold proteins, such as the Ras-Raf-MEK-ERK network (18). Further investigation of the assembly of scaffold-based signaling complexes will likely prove key to our attempts to understand and modify signaling systems within cells.

## Materials and Methods

We performed stochastic simulations as well as causality analysis using KaSim and the Kappa rule-based modeling language, and we employed the BioNetGen software package for deterministic simulations (19-21). Stochastic simulations were run until reaching an empirically determined steady-state or 10^5^ seconds in simulation-time, due to the computationally intensive nature of exact agent-based Doob-Gillespie numerical simulations (22). Both xmgrace (2D plots) and matplotlib in Python (3D plots with linear interpolation) were used for data visualization, and custom analytical tools were developed in Python (available upon request).

## Results

### Model construction

As a brief recap of the ensemble and machine signaling paradigms (described in ref. (14)), in our model of ensemble-based signaling, signal transduction events depend purely on local interactions; kinases bind the scaffold independently of one another, and a fully assembled scaffold complex need not be formed for the signal to propagate (Fig. 1A, black and red lines) (14, 23). In contrast to ensemble-like signaling, proteins associate with the scaffold in a particular order in our machine-like signaling models, so that the possible binding reactions are driven by the global state of the complex. In this paradigm, signaling machines are constructed in a hierarchical manner (Fig. 1A, red lines), ultimately forming a multi-subunit enzyme that activates downstream components (*e.g*. transcription factors) only when fully formed (14). Finally, the solution model operates as a set of independent kinases that directly bind and phosphorylate the next protein in the cascade (24).

**Figure 1.**
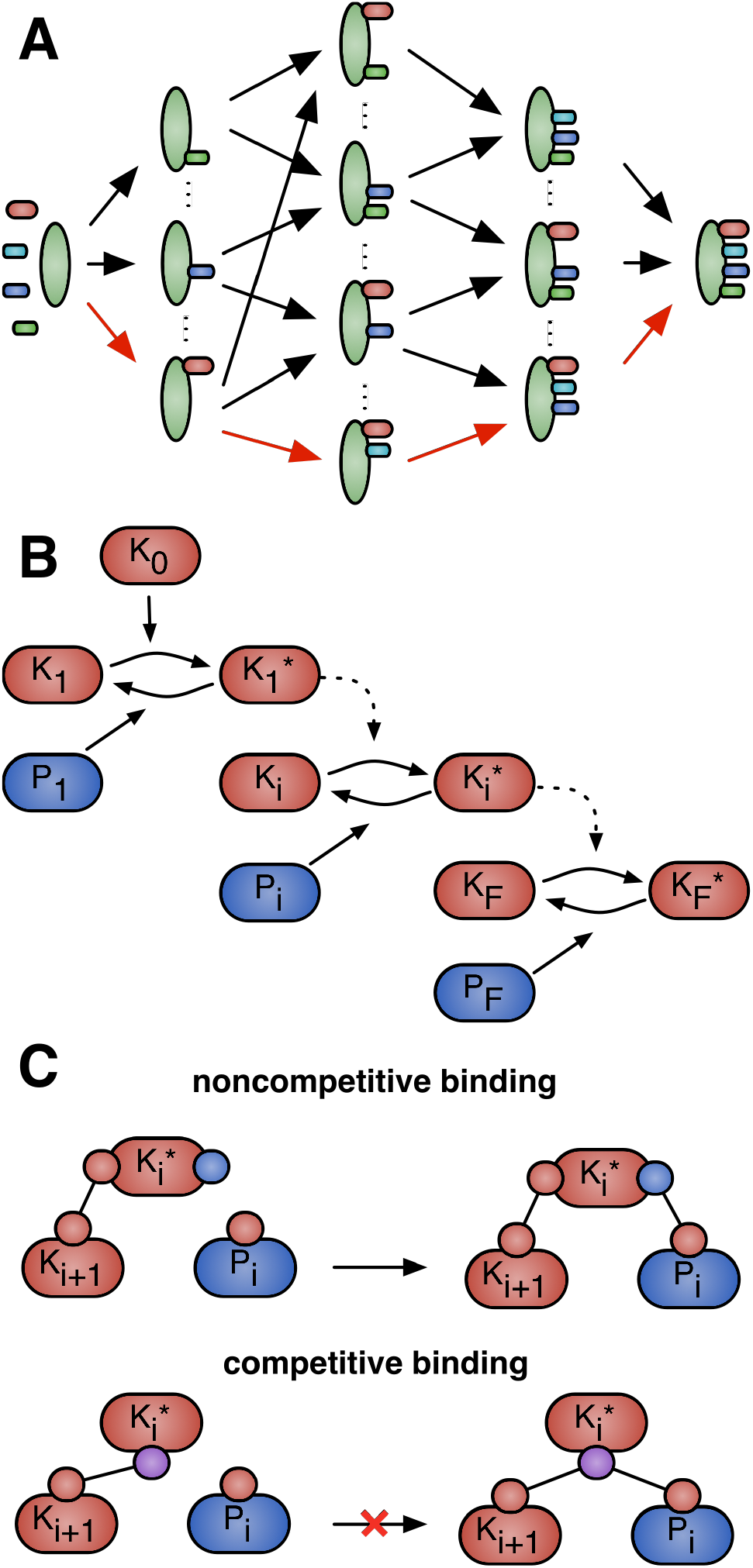
Schematics of key interaction types in scaffold-dependent signaling paradigms. **(A)** Signaling components, e.g. kinases (small, variously colored), bind to a scaffold (large, green) in order to propagate signal. Components may either bind independently of the scaffold’s binding state (black and red lines) or hierarchically (red lines), representing the ensemble signaling paradigm and machine signaling paradigm, respectively. (14, 23) Note that only in the machine signaling paradigm is the right-most complex required; ensemble signaling requires only neighboring components to be simultaneously scaffold-bound for signal propagation. **(B)** A multi-step kinase cascade based on Goldbeter and Koshland’s covalent modification cycle (26). Here, some kinase (*K_i_*) activates the next kinase in the cascade (*K_i_*_+1_ → *K_i_*_+1_^*^) and its associated phosphatase (*P_i_*) similarly deactivates it (*K_i_*^*^ → *K_i_*). The amount of active final kinase (*K_F_*^*^) is considered the output of the cascade. In cascades with scaffold proteins, the general mechanism remains the same, though the kinases actively engaged in the activation step must be bound to the scaffold. **(C)** Traditional enzyme kinetics involves competition in binding between a kinase’s phosphatase and substrate (*competitive* binding, bottom). However, since the machine and ensemble signaling paradigms allow phosphatases to bind and dephosphorylate kinases both on and off the scaffold, we implemented *noncompetitive* binding behavior (top) in the solution model as a more relevant control. Kinases in the solution models may therefore bind a substrate and phosphatase simultaneously.

The most fundamental aspect of these models’ construction is the implementation of the scaffold-kinase binding rules. For all scaffold-based models, we required that signal transduction occurs via scaffold-bound signaling species, compared to prior theoretical investigations in which the signal could propagate regardless of whether the kinases were bound to the scaffold (8, 13). Our models’ scaffold proteins are based on those found in the yeast pheromone MAPK network (25), and thus activation of any kinase in the MAPK cascade cannot occur in the absence of scaffold proteins. Furthermore, since the simulations take place in a well-mixed environment (22), our analyses are solely concerned with how the multivalent nature of scaffolds as adaptor proteins influence the dynamics of signaling and not with any spatial effects that scaffolds might have.

Stimulation of both ensemble- and machine-like cascades takes place via a signaling agent that enzymatically activates the first of a series of *N* kinases. The strength of activation is determined by modifying the catalytic rate of the first kinase’s phosphatase; signal strength is measured as the ratio of the maximum velocities of this first “stimulation” agent and the phosphatase that acts on the first kinase in the cascade (see the Supplement for further details) (26). Each subsequent kinase, which also has a corresponding phosphatase to prevent undue cascade saturation (24), binds the scaffold and propagates signal according to paradigm-specific rules. Our ensemble-like signaling models require only that an active kinase and its substrate are simultaneously bound to the scaffold for phosphorylation to occur; machine-like signaling requires that all upstream association and phosphorylation events have also occurred (Fig. 1A). The steady-state concentration of activated final kinase (*K_F_*^*^) is considered the output of the cascade, consistent with previous theoretical studies output (8, 9). Previous models of scaffold-based signaling have considered how phosphatase-based dephosphorylation of scaffold-bound kinases is implemented, and found that, in many cases, changes in qualitative signaling behavior are minimal (8, 9). Due to a lack of evidence to the contrary, we therefore assume that phosphatases may operate on scaffold-bound kinases with the same activity and parameters as freely diffusing (i.e. unbound) kinases.

In addition to our two scaffold-based signaling paradigms, we implemented a scaffoldless or *solution* model to serve as a control. This multi-stage cascade is based on the covalent modification cycle that was first mathematically characterized by Goldbeter & Koshland over 30 years ago (Fig. 1B) (24, 26). Importantly, we modified the typical representation of this process to allow phosphatase-mediated deactivation of substrate-bound kinases. This change reflects the ability of phosphatases in the machine and ensemble paradigms to dephosphorylate scaffold-bound kinases, and thus serves as an additional measure of control for the two scaffold-based models. We label this type of model *noncompetitive* since substrate and phosphatase can simultaneously bind an active kinase (Fig. 1C, top). This has the interesting impact of causing the first phosphorylation cycle, or Goldbeter-Koshland (GK) loop, in the cascade to have the properties of an isolated GK loop since there is no sequestration of the modified substrate (i.e.the second kinase) in the subsequent GK loop’s kinase-substrate complex (the *competitive* model; Fig. 1C, bottom). Said another way, the phosphatase of the initial loop may bind the kinase-substrate complex of the second loop and thus has access to the entire pool of active second kinase.

The kinetic parameters for these models were chosen based on the parameters from our previous model of the yeast pheromone signaling pathway (14). For simplicity’s sake, kinase copy numbers are identical to one another except for the final kinase, which is at a copy number that is 10-fold larger than all other kinases, and interactions between specific protein types (e.g. kinase-scaffold or phosphatase-kinase) have identical kinetics. Since the initial rate parameters we chose resulted in enzymes that were universally unsaturated (i.e. substrates were always at low concentrations compared to the *K_M_*’s of their kinases and phosphatases), we constructed a second set of parameters to consider the influence of enzyme saturation. In these saturated models, the *K_M_* of any arbitrary kinase-substrate pair was at least 2 orders of magnitude smaller than the substrate concentration (Table 1). Our results focus mainly on the models acting in the unsaturated parameter regime, since the noncompetitive nature of the phosphatases induces a strong switch-like behavior in the saturated cascades across all three signaling paradigms (see Supplement).

**Table 1.** Parameters used in our simulations. Note that all possible parameter combinations were not necessarily explored in this work. The stochastic simulation algorithm we used requires parameters to be in units relative to the number of molecules in the system (*e.g*. molecule^-1^ s^-1^ instead of M^-1^ s^-1^). In the case of association constants, conversion to units of concentration requires multiplication by a volume and Avogadro’s number. For example, given a yeast cell with a volume of 40 fL, the unsaturated *k_on_* of 10^-5^ molecule^-1^ s^-1^ is approximately 2 x 10^5^ M^-1^ s^-1^. We vary *k_on_* (the denominator of the Michaelis constant) to control the saturation of the kinases, rather than saturating the kinases by increasing the copy number of the substrates. This allows us to simulate a saturated condition without increasing copy numbers, which could alter the noise properties of the system. Note that increasing the stochastic *k_on_* corresponds to increasing substrate concentration by simulating the same number of molecules in a smaller effective volume. Abbreviations: Ensemble (e), Machine (m), Solution (s), Depth in cascade (*i*), Michaelis constant (*K_M_*).

| Parameter | Values |
| --- | --- |
| Signal range $\left(\frac{V_{max,K_1}}{V_{max,P_1}}\right)$ | $10^{-5}$ to $10^5$ by $10^{1/5}$ (e, s)<br>$10^{-8}$ to $10^2$ by $10^{1/5}$ (m) |
| Phosphatases (molecules, $i > 1$) | 50 to 500 by 50, 750, 1000 (all) |
| Scaffolds (molecules) | 10, 100, 1000, 2000, 5000, 10000, 20000, 40000 (e, m) |
| Kinases (molecules) | 1000 (all) |
| $K_M$ (molecules) | $\sim 10^5$ (unsaturated, $k_{on} = 1 \times 10^{-5}$ molecule$^{-1}$ s$^{-1}$)<br>$\sim 1$ (saturated, $k_{on} = 1$ molecule$^{-1}$ s$^{-1}$) |

### Steady state dose-response trends

The first step in characterizing these signaling paradigms was to generate sets of dose-response data while varying key aspects of the cascade, namely the phosphatase copy number and the number of distinct kinase types in the cascade (equivalent to the number of kinase binding sites on the scaffold, which we refer to as the cascade’s *depth*; Fig. 2A). The resultant dose-response trends were universally sigmoidal in shape, and so we characterized the behavior of each model by fitting the response data to a Hill function:

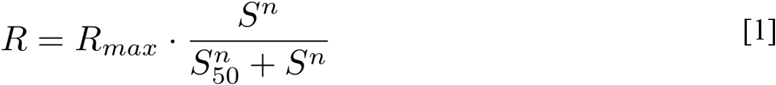

**Figure 2.**
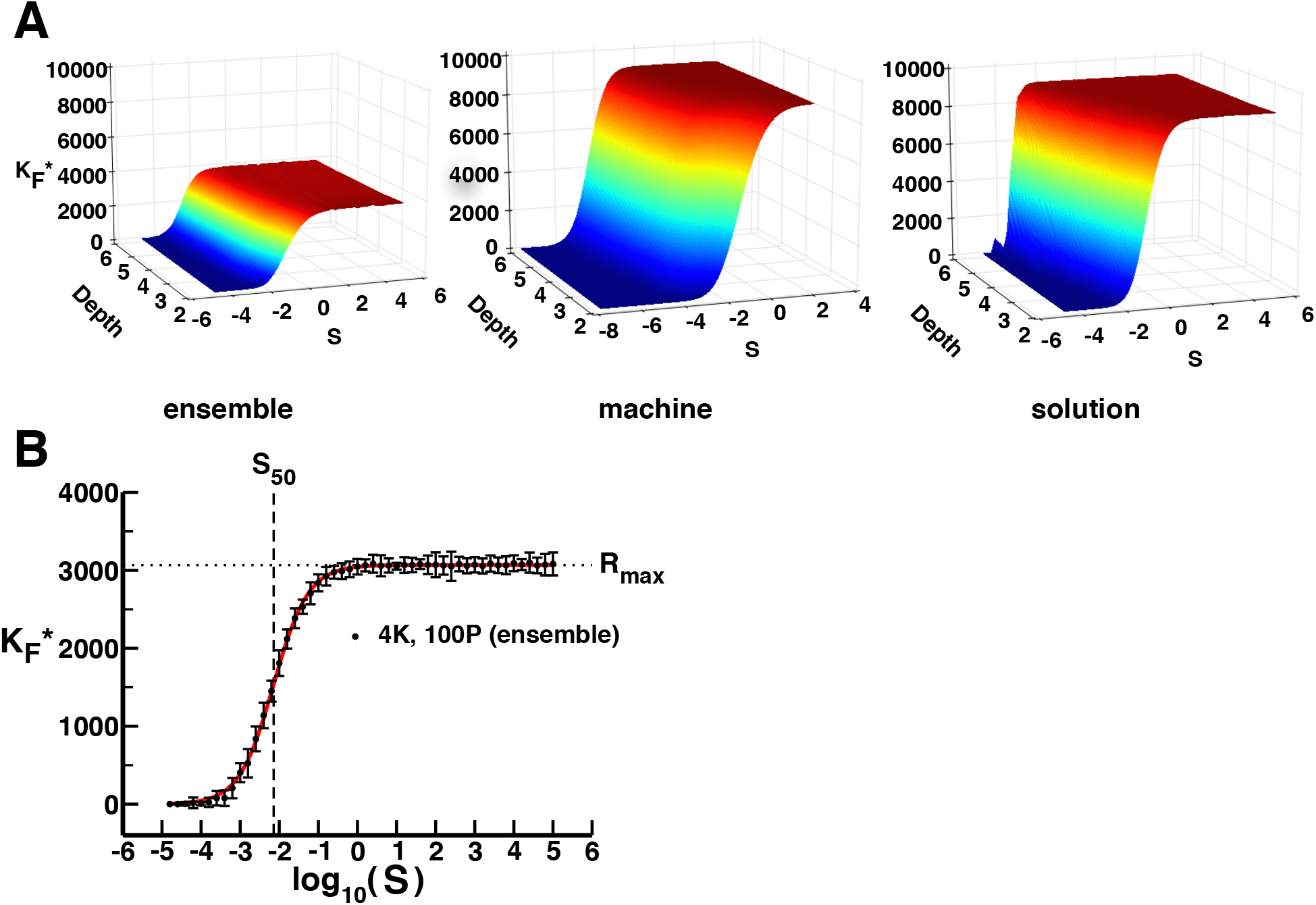
Dose-response dynamics for the different signaling paradigms. **(A)** Dose-response surfaces for unsaturated, low phosphatase simulations of the three signaling paradigms. Depth describes the number of stages in the multi-kinase cascade, *S* is signal strength, and *K_F_*^*^ is the number of active final kinases, which we consider to be the output. We used simple linear interpolation to smooth these surfaces. **(B)** A representative data set from ensemble model simulations (10 replications, with 95% confidence intervals about the mean) for a depth of 4 kinases (4K) and a 1:10 phosphatase to kinase ratio (100P). The x- and y-axes (*S* and *K_F_*^*^, respectively) are as in (A). The solid line is the 3-parameter Hill function fit, where *R_max_* = 3067 (dotted line), *S_50_* = 0.00717 (dashed line), and *n* = 0.991; all parameters are statistically significant (*p* < 10^−16^).

We can thus describe the steady state response properties of each model in terms of the Hill function’s parameters: maximum response (*R_max_*), sensitivity to signal (*S_50_*; the signal producing a half-maximal response), and response ultrasensitivity (*n*; the sharpness of the switch from minimum to maximum response). A representative data set is shown in Fig. 2B with the *R_max_* and *S_50_* parameters obtained from the fit indicated. In general, our analyses refer mainly to models with 100 phosphatases for each kinase, so that there is a 1:10 ratio of phosphatases to kinases (except with the final kinase in the cascade where there is a 1:100 ratio) unless otherwise noted. This allows for stronger signal throughput as compared to models with higher phosphatase to kinase ratios.

The Hill parameter governing maximal response, *R_max_*, is nearly identical between machine and solution models when both are in the same parameter regime (Fig. 2A). For these two paradigms, over 90% of final kinase pool is active at steady state when stimulated with a strong activating signal, regardless of whether the kinases in the cascade are unsaturated or saturated. On the other hand, the unsaturated ensemble models exhibit a much lower maximum response, with about 40% activation of the final kinase concentration even at very high levels of cascade stimulation. This indicates that the maximum response of a network is much more dependent on the rules governing the protein interactions than the presence of a scaffold protein, a trend that is consistent throughout this work. In other words, it is not the mere presence of the scaffold itself, but rather how the scaffold-based signaling complex assembles that ultimately determines *R_max_*.

Prior theoretical studies have shown that increasing the depth of a scaffoldless cascade increases the sensitivity to signal (i.e. decreases *S_50_*) (24, 26). Our results support this claim (Fig. 3A), despite operating in a different parameter regime. Similar to the maximal response trends described above, addition of a scaffold protein alters the quantitative response in a paradigm-dependent manner. The ensemble models exhibit increased sensitivity to signal as a function of cascade depth, but the increase is shallower than the increase observed for solution models. The increase in sensitivity with cascade depth for the machine models, on the other hand, is much sharper, further highlighting the fact that the knowledge of the binding mechanisms between kinases and scaffold proteins is central to understanding how scaffolds perturb the response to incoming signals. As a side note, the sensitivity of saturated scaffold-based simulations is essentially invariant with respect to cascade depth when the phosphatase-to-kinase ratio is 1:10 (see Supplement).

**Figure 3.**
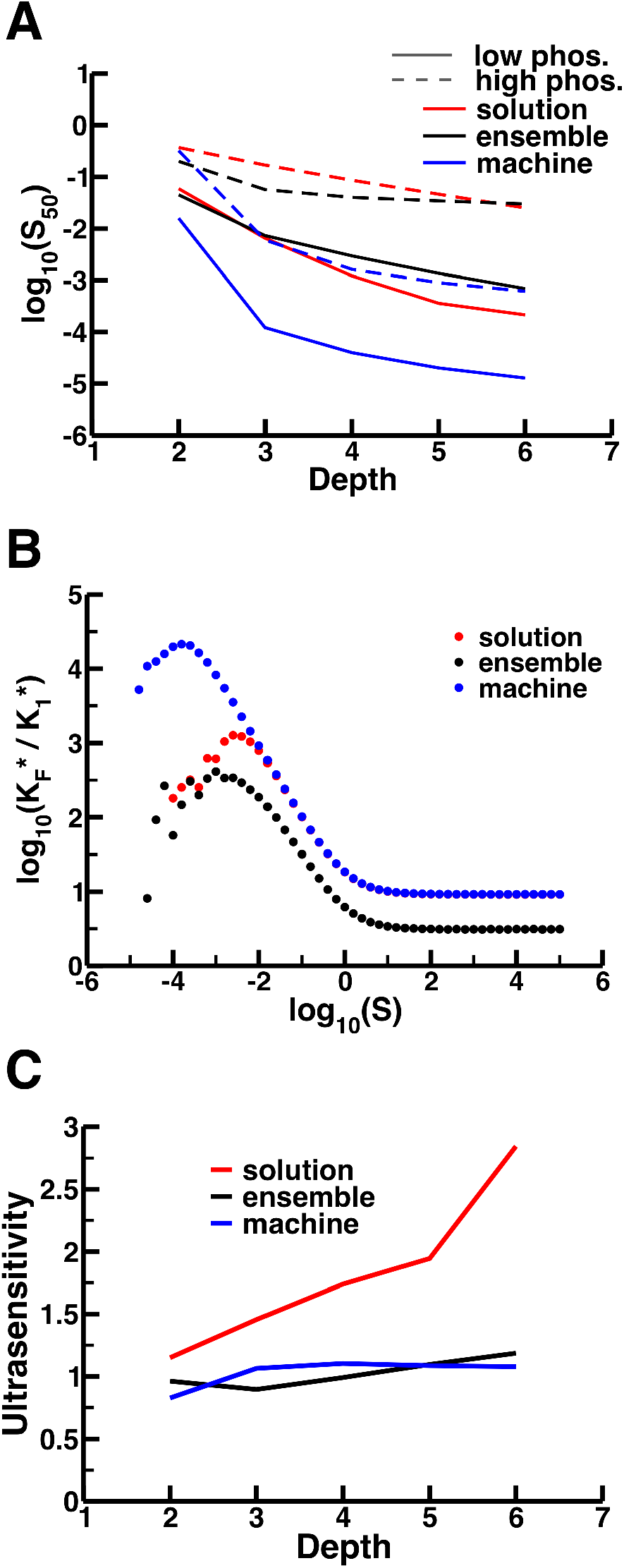
Scaffolding generates both general and paradigm-specific behaviors **(A)** Examining sensitivity to signal (y-axis) given the depth of the cascade (x-axis) for both high (dashed lines, *P_i_* _> 1_ = 1000) and low (solid lines, *P_i_* _> 1_ = 100) phosphatase activity reveals that lower phosphatase activity, as well as increased cascade depth, leads to increased sensitivity. Notably, for cascades with depth ≥ 3, machine-based signaling generally exhibits increased sensitivity to signal compared to ensemble and solution based signaling regardless of the level of phosphatase activity. **(B)** Signal amplification, defined as the ratio of first to last kinase activity in a cascade (*K_F_^*^* / *K_1_^*^*), occurs in all signaling paradigms. The data shown here are taken from models with depth = 4. The underlying cause is a shift in signal sensitivity with cascade depth (A), which induces this amplification at moderately low signal levels. **(C)** Scaffolding decreases the ultrasensitivity of the response in unsaturated models with low phosphatase activity. Despite the stark difference in scaffold protein assembly in the ensemble and machine paradigms, the scaffold has a similar “linearizing” effect relative to the solution paradigm.

This increase in sensitivity to signal with cascade depth directly impacts another posited role in signaling dynamics for scaffolds, which is a mechanism for prevention of signal amplification (3, 4, 8). Specifically, the hypothesis is that scaffold proteins might limit signal amplification due to stoichiometric constraints on the assembly of relevant signaling species. In order to examine this systematically in our three signaling paradigms, we defined signal amplification similarly to Locasale, *et al.* as the ratio of the final kinase’s activity to the first kinase’s activity: *K_F_^*^* / *K_1_^*^*. Our results reveal that all signaling paradigms exhibit some degree of signal amplification at moderately low levels of signal (Fig. 3B, Supplement). The reason for this, as alluded to above, results from the increased sensitivity corresponding to increased cascade depth (Fig. 3A). As the depth of a cascade increases the relatively low signal levels that activate only a small portion of the *K_1_* pool (which behaves as a substrate within an isolated GK loop in all three signaling paradigms) may subsequently activate all final kinase molecules (3, 4, 8).

Additionally, the presence of modified scaffold proteins in a signaling network has been shown to modify the steepness of the dose-response curve (7). As a result, we expect that differences in scaffold implementation could impact the steepness or ultrasensitivity of the dose-response curve as characterized by the Hill coefficient, *n*. Our models show reduced ultrasensitivity for the ensemble and machine paradigms as compared to the solution paradigm in the unsaturated, low phosphatase parameter regime (Fig. 3C). Quantitatively, the saturated cascades have a much larger *n* relative to the unsaturated cascades (as might be expected from prior analyses of GK loops (24, 26)). It is important to note that our simulated data sets lack the signal-space resolution for accurate characterization of the Hill coefficient, *n*, for saturated models, since they are all extremely ultrasensitive compared to the unsaturated models. Further simulations would need to be performed to thoroughly characterize the extreme ultrasensitivity of the saturated models, and this computationally expensive task is outside the scope of this study. Nonetheless, previous hypotheses regarding scaffold-induced dose-response linearization are supported by our findings for both the machine and ensemble models in the unsaturated regime.

### Speed and reliability of response

In addition to steady state dose-response behavior, other properties of signaling networks could easily contribute to their function and evolution. One such property is the speed at which cells are able to respond to some environmental stimulus. We explored the influence that scaffold proteins have on the speed of response by calculating the time it takes for a simulation to reach a response greater than half of that observed at steady state (*T_50_*). We calculated this value at two signaling strengths: the signal nearest that required to reach half-maximal response (*S_50_*) and the signal resulting in maximal response (*S_max_*). In the unsaturated models, *T_50_* increases monotonically with cascade depth for all three signaling paradigms at both *S_max_* and *S_50_* (Figs. 4A and 4B). However, the machine model consistently takes longer to respond, likely due to the time required to successfully assemble discrete signaling machines on the scaffold. In fact, the machine model does not reach *T_50_* for nearly one day of simulated time for signal values nearest *S_50_* (Fig. 4A). Note that the association rates employed here (>10^5^ M^−1^ s^−1^ assuming a volume similar to that of a yeast cell, Table 1) represent fairly fast binding kinetics for proteins, so the relatively slow response in this case is not due to unrealistically slow kinetic rates. On the other hand, there is negligible difference between the ensemble and solution model response times at both signal values (with the exception of the two-kinase cascades). As observed above, the influence of a scaffold on signaling dynamics in this case is highly dependent on the nature of the binding rules themselves.

**Figure 4.**
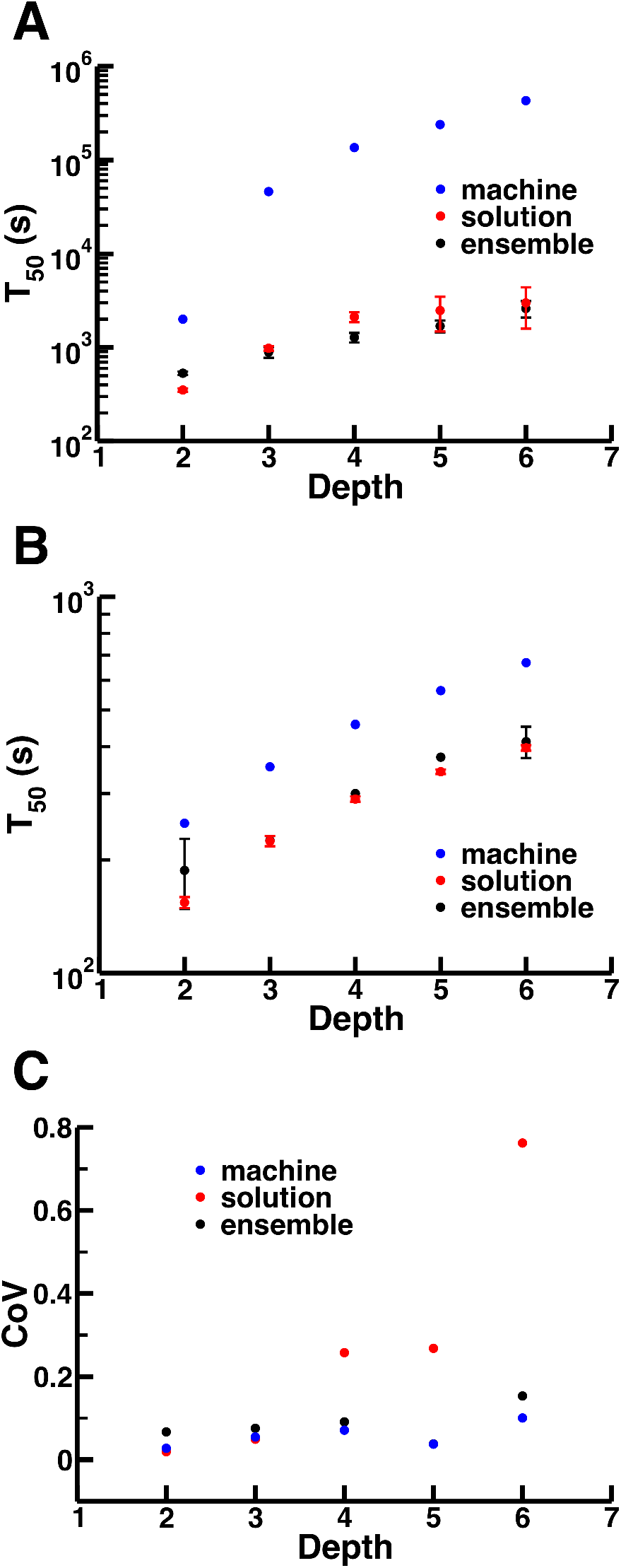
Scaffolding alters variability and speed of response. **(A & B)** Response time at *S_50_* (A) and *S_max_* (B) increases with cascade depth. These plots indicate the length of time (in simulated seconds) taken to exceed half the observed response at steady state (*T*_50_; y-axis) as a function of cascade depth (x-axis). Similar to *R_max_* and *S*_50_ (Figs. 2A and 3A, respectively), the response time does not specifically depend on the presence of the scaffold, but rather on the assembly paradigm. Strikingly, the machine model exhibits response times nearly two orders of magnitude greater than those observed in ensemble and solution models for deeper cascades with intermediate signal strength (A). **(C)** Scaffold proteins suppress the noise present in deep solution cascades independent of the assembly paradigm. The coefficients of variation (y-axis) are taken from the simulation whose signal is nearest the fitted *S_50_* value.

In addition to providing a timely response, some signaling networks must reliably respond to signals on the single-cell level (e.g. gradient tracking for chemotaxis or shmoo formation in yeast) (27). Reduction of biochemical noise may thus be a key property of signaling cascades, and we posit that scaffold proteins could play a role in controlling fluctuations. To test this possibility, we examined the variability in response (as measured by the coefficient of variation) for simulations with signal values nearest to their respective *S_50_* values and found that scaffolds strongly reduce intrinsic noise for intermediate response values (i.e. the steepest region of the dose-response curve), especially for relatively deep cascades (Fig. 4C). This makes intuitive sense in the case of the machine model: by constructing a discrete multimeric enzyme (signaling machine) instead of relying on a series of GK loops to activate the final kinase in a cascade, the machine model exhibits less variability in active kinase numbers during signal transduction, thus limiting noise. Perhaps more interesting, however, is the fact that the ensemble models also significantly reduce noise levels, which is particularly striking when considering that the ensemble models are sufficiently combinatorially complex to generate nearly an order of magnitude more signaling species than the machine and solution models in deeper cascades (Supplement). These results indicate that scaffold proteins provide a mechanism for reducing fluctuations regardless of how they assemble, damping the noise that can arise from the strong response amplification present in cascades that do not utilize a scaffold.

### Effects of scaffold number variation

The results described above were all produced with a stoichiometric ratio of scaffold proteins to kinases for kinases 1 through *N* – 1 (where *N* is the cascade depth). It has been shown repeatedly that variations in scaffold concentration can have strong and sometimes nonintuitive effects on network response. A common example is the *prozone* effect, or *combinatorial inhibition* (9, 12). We therefore examined our models in the context of scaffold copy number, varying this quantity by over two orders of magnitude.

As previously studied in a model of the yeast pheromone MAPK network, machine-based signaling does not exhibit the experimentally-verified effect of combinatorial inhibition (14). Instead, the machine model exhibits an upper limit on *R_max_* that is realized near stoichiometric copy numbers and *R_max_* does not decrease as the concentration of scaffold increases (Fig. 5A, right). This is similar to previous results where increasing cooperativity between the scaffold’s binding partners results in a decrease in combinatorial inhibition (13). The ensemble models, however, do exhibit combinatorial inhibition, with peak *R_max_* near stoichiometric scaffold concentration that drops sharply at higher scaffold concentration (Fig. 5A, left). This decrease becomes more pronounced as the cascade depth becomes larger: as the valency of the scaffold increases, so does the combinatorial complexity of the system, and thus the influence of combinatorial inhibition is larger.

**Figure 5.**
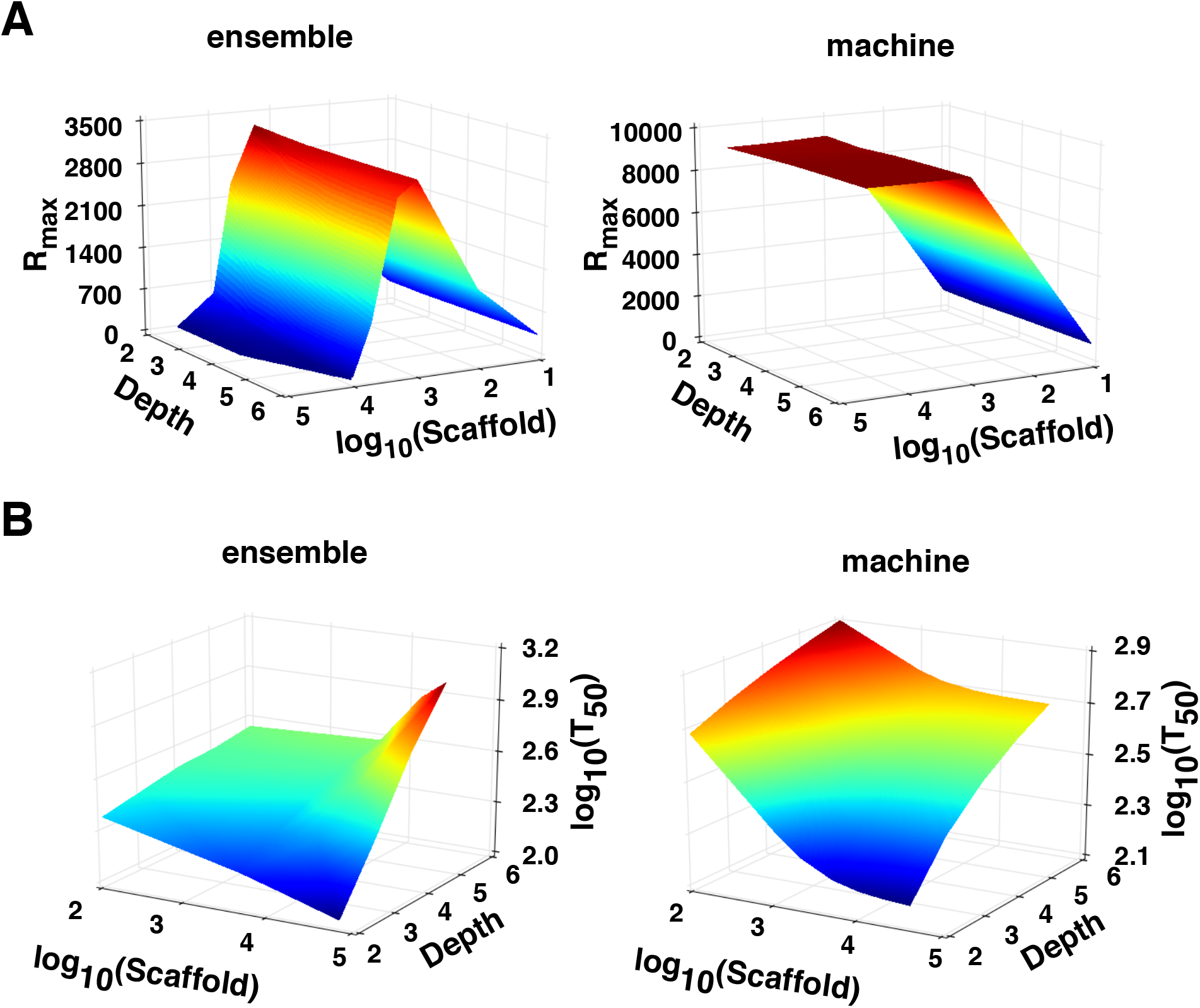
Scaffold concentration modulates select signaling behaviors. **(A)** Maximum response as a function of depth and scaffold number. Consistent with findings from prior experimental and theoretical studies (9, 12), we observe combinatorial inhibition due to high concentrations of scaffold proteins in the ensemble model simulations (left). Contrary to this, the machine model produces no such inhibitory effect since the hierarchical nature of signaling machine assembly prevents the combinatorial explosion of scaffold-based species that is present in the ensemble model (14). **(B)** Response time as a function of depth and scaffold number at *S_max_*. We observe a universal decrease in response time with respect to scaffold number in the machine model for all cascade depths (right), though in the parameter space considered here, this increase is less than an order of magnitude. Increasing scaffold numbers in the ensemble model, while showing faster responses in the 2-kinase cascade (likely due to the relative similarity between the 2-kinase ensemble model and the 2-kinase machine model rule structures, in addition to the reduced response due to combinatorial inhibition), displays slower response times for deeper cascades as a result of increased combinatorial complexity.

We also observed that both assembly paradigms are less sensitive to signal (i.e. their *S_50_* increases) when the scaffold copy number is decreased for a cascade depth of 3 (Figure S6). This is consistent with experimental findings for the KSR1 scaffold involved in the mammalian Ras-Raf-MEK-ERK MAPK cascade (28). Increasing scaffold number results in increased sensitivity to signal up to some saturating maximal sensitivity in both models. The maximum sensitivity observed in machine models is due to the fact that, once the scaffold reaches the same copy number as the kinases in the cascade, additional scaffold molecules do not bind kinases and thus have a minimal effect on steady-state response (Fig. 5A, right). The saturation in sensitivity for the ensemble models occurs because, at high scaffold concentrations, combinatorial inhibition essentially prevents any response at all (see Supplement). We found that varying scaffold numbers in the machine and ensemble signaling paradigm has little noticeable effect on ultrasensitivity or the noise in response (see Supplement).

Another signaling property that shows a clear scaffold-dependent trend is the variation of *T_50_* with scaffold concentration. In the machine models, increased scaffold numbers universally decrease the measured *T_50_* over the explored range of cascade depths until reaching a limit between 5000-10000 scaffold proteins (Fig. 5B, right). This occurs because higher scaffold concentrations raise the probability of initiating machine assembly, increasing the frequency of association between the scaffold and the first kinase in the cascade. This phenomenon is also present in the 2-kinase ensemble model, possibly due to the fact that the model essentially builds a 2-subunit signaling machine (though the reduced *R_max_* resulting from combinatorial inhibition may also contribute to lower *T_50_* values; Fig. 5B, left). For deeper ensemble cascades, increasing the scaffold copy number raises the *T_50_* as does the presence of combinatorial inhibition: as scaffold numbers grow, the time it takes to propagate signal also grows due to sequestration of signaling components on different scaffold molecules.

### Crosstalk

The ubiquity of crosstalk between signaling pathways (defined as one pathway’s signaling components influencing another pathway’s activity) in eukaryotic organisms is uncontested (29, 30), and it is currently unclear how any degree of specificity is maintained in the face of this abundant crosstalk (6, 31). One supposition is that scaffold proteins act as some sort of intracellular circuit board, directing signal transduction towards specific outputs for any given input (3). It is likely that the assembly paradigm will influence the efficacy of crosstalk prevention in signaling cascades: in the absence of some sort of well-defined signaling complex (i.e. machine), it is unclear how scaffolds could prevent cross-pathway activation. To examine this, we adapted our three model types to include two pathways, each with a scaffold (except the solution model) and a shared kinase, in this case, the second in a 4-kinase cascade, *K_2_* (Fig. 6A). Motivating this analysis is the existence of parallel MAPK networks in yeast that involve distinct scaffold proteins (Ste5 in the pheromone signaling network and Pbs2 in the osmolarity response pathway) but share an upstream kinase (Ste11) that binds to both scaffolds (1).

**Figure 6.**
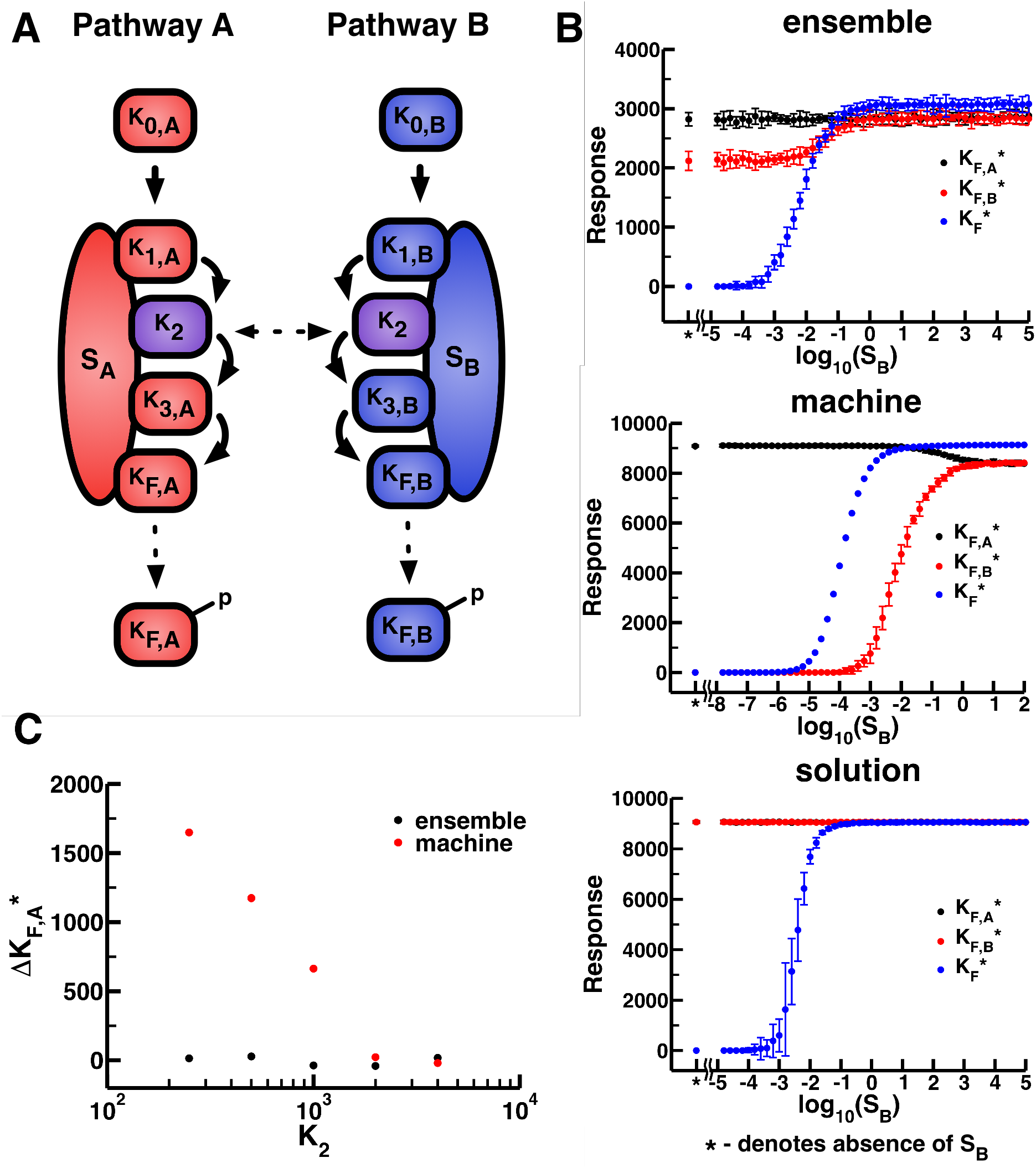
Crosstalk in the three signaling paradigms. **(A)** Schematic of crosstalk in scaffold-based signaling networks. In this figure, red components belong exclusively to pathway A while blue components belong to pathway B. The second kinase in both cascades (purple) is shared. Here, solid lines represent activation events, while dotted lines show translocation. In the solution model, kinases bind to one another, thus both *K_1,A_* and *K_1,B_* are capable of activating and binding *K_2_*, which then binds and activates both *K_3,A_* and *K_3,B_*. **(B)** Various cascade outputs (y-axis) as a function of log-transformed signals in pathway B (x-axis); pathway A is exposed to maximum signal (*S_A_* = 10^5^ for ensemble/solution simulations and *S_A_* = 10^2^ for machine simulations) for all data points. Black and red points indicate pathway A and B response, respectively. As a reference, blue points show the response for a single pathway model stimulated with *S_B_*-strength signal. **(C)** Difference in pathway A response (y-axis) between a model with maximal stimulation of pathway A and minimal stimulation of pathway B and a model with maximal stimulation of both pathways as a function of the number of shared kinases (*K_2_*; x-axis). As seen in panel (B), maximal activation of both pathways in the machine signaling paradigm introduces a decrease in output relative to maximal activation of only pathway A, whereas this difference is negligible in the ensemble model. This occurs when *K_2_* is the limiting factor in the signaling cascade (*i.e*. the component with the lowest copy number).

Initially, we examined the effects for a scenario with a maximally activated pathway A (*S_A_* = 10^5^ in the ensemble and solution models and *S_A_* = 10^2^ in the machine models), combined with a minimally activated pathway B (*S_B_* = 10^−5^ in the ensemble and solution models and *S_B_* = 10^−8^ in the machine models). We found that the solution model exhibited equal response from both pathways, despite stimulating only one (Fig. 6B, bottom). This is intuitive when considering that the 3^rd^ kinase, *K_3_*, in each pathway competes equally for the pool of active shared *K_2_*. Similarly, an active *K_2_* in the ensemble model may bind either scaffold A or scaffold B if it dissociates from pathway A’s scaffold. Despite this fact, pathway A maintains a higher steady-state response than pathway B in this scenario (Fig. 6B, top). This is likely due to the additional biochemical events necessary for *K_2_* to activate pathway B. Upon its activation, the shared kinase can immediately phosphorylate the 3^rd^ kinase in pathway A, assuming *K_3,A_* is already present on pathway A’s scaffold protein. Activation of *K_3,B_* requires an additional dissociation event (as mentioned above) and an association event (*K_2_* binding pathway B’s scaffold), presumably resulting in a lower absolute response. This can be visualized via causality analysis tools present in the KaSim software package (see Supplement) (20, 32). Finally, no inappropriate cross-pathway activation exists in the machine model in this scenario (Fig. 6B, middle) since assembly of the signaling machine requires that pathway B’s first kinase is active, which occurs very infrequently due to the low external activation of pathway B. In both the ensemble and machine models, increasing the signal input to pathway B eventually causes it to respond at levels similar to those of pathway A (Fig. 6B).

The machine model is thus the only signaling paradigm we examined that prevents one pathway from activating a second pathway where the second has no (or minimal) signaling input. However, an alternative form of crosstalk still arises in the machine paradigm and can be observed in Fig. 6B. In the case of our initial crosstalk models, the shared kinase is the limiting factor in signal transduction (i.e. the component with the smallest copy number) since its per-pathway concentration is halved. Competition for this kinase alters signal throughput, albeit in a different way than inappropriate activation of another pathway’s output. In this case, the activity of one pathway is reduced when its components are recruited to another active pathway, and the output of pathway A actually *decreases* as pathway B becomes fully active (Fig. 6B, middle). To better characterize this phenomenon, we calculated the difference in *K_F,A_*^*^ between two cases: case 1, where pathway A is maximally active and pathway B is inactive, and case 2, where both pathways are maximally active. We represent the difference between case 1 and 2 as Δ*K_F,A_*^*^. Due to the sequestration of the shared kinase, the machine model with a *K_2_* concentration of 500 molecules (i.e. half the concentration of the scaffold) has a Δ*K_F,A_*^*^ > 1000, indicating that full activation of pathway B can reduce the total pool of active *K_F,A_* by 10% (Fig. 6C). As one would expect, this drop in response output is mitigated with an increase in *K_2_* concentration: doubling the *K_2_* concentration relative to the scaffold (i.e. *K_2_* = 2000) results in essentially identical response from the first pathway regardless of the second pathway’s level of stimulation. Thus, even if separate scaffolds nucleate formation of a machine-like signaling complex in two pathways, establishing true independence between those pathways requires detailed knowledge of the relative concentrations of the scaffold and any kinase that is shared between them. Interestingly, limiting *K_2_* concentrations do not generate similar behaviors in the ensemble models (Fig. 6C), likely due to the fact that the shared kinase is not sequestered in an assembled (or assembling) signaling complex.

## Discussion

Our results clearly indicate that the dynamical features of a signaling cascade can be drastically influenced not just by the presence of a scaffold protein, but also how the kinases in that cascade assemble onto the scaffold itself. These findings are summarized in Fig. 7. Strikingly, we found that only two of the response characteristics we considered were similar between the ensemble and machine signaling paradigms. The most notable of these was the fact that presence of a scaffold protein in the cascade universally reduces noise and variability in molecular responses. Suppression of noise would clearly be advantageous in cases where individual cells must gather accurate information about their environment, such as determining if a potential mating partner is present (7, 14, 27, 33). This is likely related to the fact that scaffolds also generally linearize dose-response behavior, preventing the massive increase in ultrasensitivity that generally occurs as kinase cascades become deeper (24, 26).

**Figure 7.**
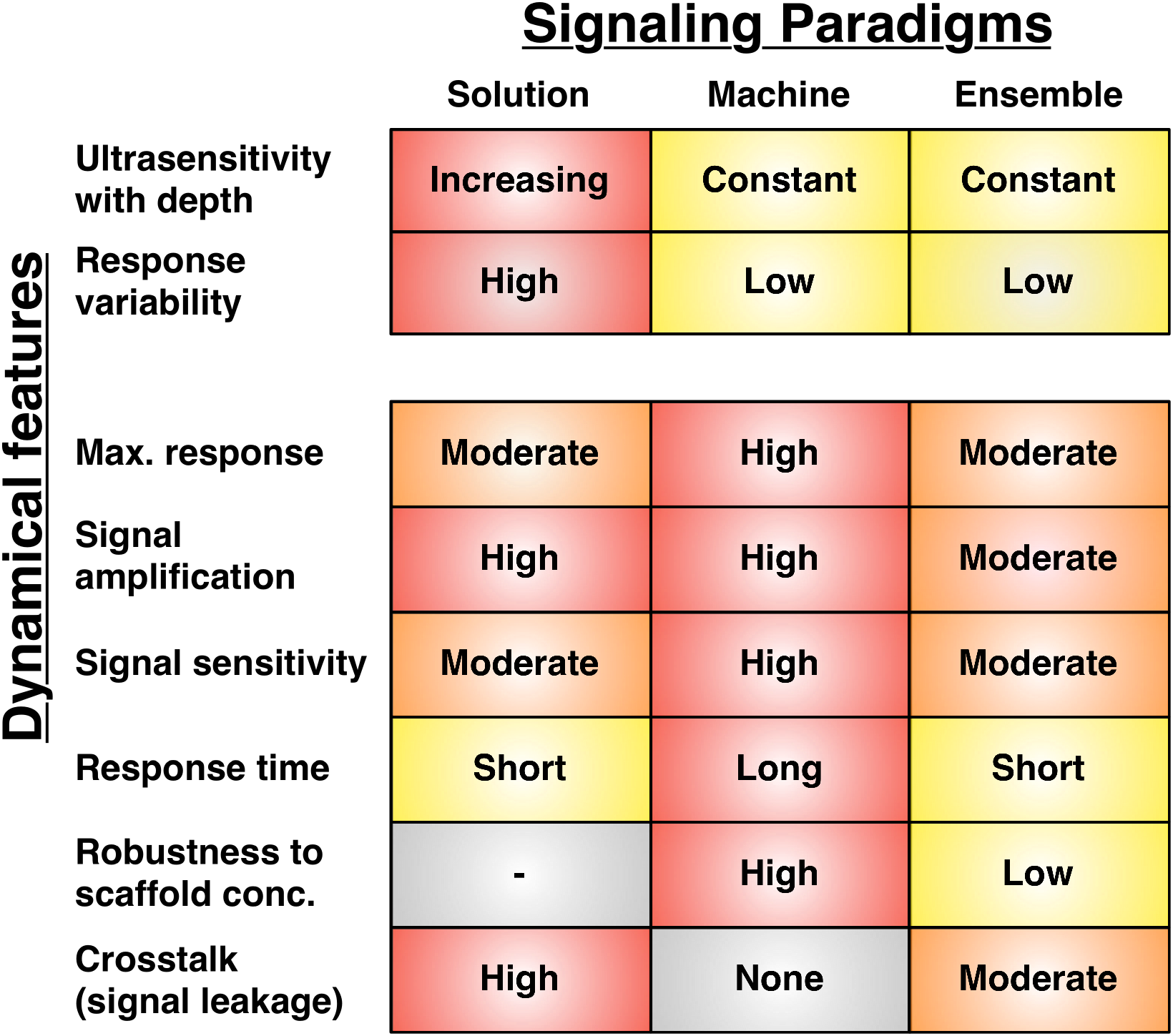
Comparison of various features between signaling paradigms. The three columns represent the three distinct types of signaling models considered in this work. Each row corresponds to a different dynamical feature. For two of these features, namely the variability of the response and the change in ultrasensitivity as cascades become deeper, the two scaffold-based signaling paradigms demonstrate similar behavior. In all other cases, however, the manner in which scaffold-based signaling complexes assemble is as important as whether or not the cascade uses a scaffold in the first place. Note that the results summarized in this figure are for unsaturated models of varying depths.

The majority of the dynamic features we considered, however, showed strong dependence on how the kinases actually assemble onto the scaffold itself (Fig. 7). Machine-like structures generate higher absolute levels of output than ensembles, but require much longer times to achieve those responses, especially when signals are near the half-maximal level. Ensembles, on the other hand, can exhibit high degrees of combinatorial inhibition if scaffold concentrations are not tightly maintained near stoichiometric concentrations. The two assembly paradigms also have very distinct behaviors in terms of how they handle components that are shared between multiple pathways. The machine model exhibits complete insulation from inappropriate activation by other pathways with shared downstream components, while the ensemble model does not. However, our models predict that scaffold proteins will reduce cross-pathway activation even in the ensemble case, improving signaling specificity relative to cascades that have no scaffold at all. These specific predictions can be used to inform experiments in the yeast MAPK network, upon which our models are based. In particular, the shared kinase Ste11 binds to scaffolds in both the pheromone response pathway (Ste5) and the high osmolarity response pathway (Pbs2). From the results seen in Figure 6, maximal activation of both pathways should not affect the relative pheromone response as compared to the response given maximal activation of the pheromone pathway with no activity in the high osmolarity pathway, regardless of the quantity of Ste11 in the system if the Ste5 scaffold binds its effectors independently. This should be relatively easy to test experimentally, requiring only a mechanism to vary the expression of Ste11 and a reliable means to independently stimulate the two pathways.

In general, these observations suggest that various assembly paradigms could play strikingly different evolutionary roles (3). The nature of the signaling machine’s complex structure and hierarchical assembly is reminiscent of highly-conserved multi-subunit proteins like the ribosome (34). In this paradigm, the lack of combinatorial inhibition and decreased signaling time with increases in scaffold concentration indicate a resistance to fluctuations in protein concentration, which might arise due to the inherent noise in gene expression or from other, possibly “extrinsic,” sources (35). These traits couple well with scaffold-specific, but paradigm-independent, properties, such as reduced dose-response ultrasensitivity and noise in response (Figs. 3, 4). Scaffold complexes that assemble like machines can thus provide finely tuned and phenotypically robust behaviors. However, this type of multi-subunit protein might be difficult to evolve in comparison to the ensemble paradigm, since the scaffold would need to evolve extensive allosteric communication among its subunits in order to enforce hierarchical assembly (e.g., the fact that kinase *i* will not bind the scaffold until kinase *i* - 1 is already present in the complex, Fig. 1). Adding a new kinase to the cascade, or generating an entire signaling machine *de novo*, would thus likely require a rather lengthy process of evolving those constraints. In contrast, adding a new kinase to the ensemble model simply involves adding the relevant binding domain somewhere in the scaffold. Interestingly, extensive experimental work has shown that Ste5, the prototypical MAPK scaffold, can easily accommodate this kind of novel interaction, often generating highly functional dynamics just by adding new interactions or shuffling existing ones (7, 36, 37). Ensembles thus exhibit a much higher degree of functional plasticity, generating *weak regulatory linkage* among signaling components and enabling the rapid evolution of new phenotypes (38). Weak linkage, coupled with other ensemble-specific features (e.g. noise suppression, fast responses to signal) could provide strong fitness advantages in rapidly changing environments. In essence, scaffold proteins in the ensemble paradigm facilitate the evolution of additional cellular functionality (e.g. “rewiring” signaling pathways) whereas scaffold proteins in the machine paradigm better conserve existing cellular functions (e.g. reliably constructing ribosomes) (38).

While our findings indicate that the specific mechanisms of scaffold complex assembly are important for the function and evolution of signaling networks, little is known empirically about the process itself in living cells. Since Ste5 exhibits combinatorial inhibition, and is quite tolerant to the addition of new interactions or the permutation of existing ones, it is fairly likely that Ste5 signaling exhibits at least some ensemble-like properties (7, 14, 36, 37). While there is some recent work examining the assembly of other multivalent scaffolds *in vitro* (39), the generality of the ensemble paradigm is currently unclear. Our work suggests that several relatively simple experiments could be helpful in establishing whether or not a particular scaffold assembles as a machine or an ensemble. For instance, varying scaffold concentrations by under- and over-expressing the protein and measuring the variation in response speed and steady-state response level could provide at least some preliminary indication of the assembly pathway involved (e.g. Fig. 5). Mutations aimed at disrupting allosteric communication among subunits (e.g. by replacing wild-type interaction domains with novel ones) could also be helpful in assaying the assembly paradigm employed by any given scaffold.

Scaffold proteins have been recognized both as important drug targets (such as the Kinase Suppressor of Ras (KSR) in the MAPK/ERK cascade) and as key components in the design of synthetic or biologically-inspired signaling systems (3). Our work indicates that any attempt to rationally control the behavior of a scaffold-based signaling cascade, either through small molecules or through engineered mutations, must consider how the complexes themselves are assembled. Experimentally characterizing these assembly processes for a wide range of scaffold proteins thus represents a key unmet challenge in systems and synthetic biology.

## References

1. Chen RE & Thorner J (2007) Function and regulation in MAPK signaling pathways: lessons learned from the yeast Saccharomyces cerevisiae. Biochim. Biophys. Acta 1773(8):1311-1340.

2. Chen WW, et al. (2009) Input-output behavior of ErbB signaling pathways as revealed by a mass action model trained against dynamic data. Mol. Syst. Biol. 5:239.

3. Good MC, Zalatan JG, & Lim WA (2011) Scaffold proteins: hubs for controlling the flow of cellular information. Science 332(6030):680-686.

4. Burack WR & Shaw AS (2000) Signal transduction: hanging on a scaffold. Curr. Opin. Cell Biol. 12(2):211-216.

5. Patterson JC, Klimenko ES, & Thorner J (2010) Single-cell analysis reveals that insulation maintains signaling specificity between two yeast MAPK pathways with common components. Sci. Signal. 3(144):ra75.

6. McClean MN, Mody A, Broach JR, & Ramanathan S (2007) Cross-talk and decision making in MAP kinase pathways. Nat. Genet. 39(3):409-414.

7. Bashor CJ, Helman NC, Yan S, & Lim WA (2008) Using Engineered Scaffold Interactions to Reshape MAP Kinase Pathway Signaling Dynamics. Science 319(5869):1539-1543.

8. Locasale JW, Shaw AS, & Chakraborty AK (2007) Scaffold proteins confer diverse regulatory properties to protein kinase cascades. Proc. Natl. Acad. Sci. USA 104(33):13307-13312.

9. Levchenko A, Bruck J, & Sternberg PW (2000) Scaffold proteins may biphasically affect the levels of mitogen-activated protein kinase signaling and reduce its threshold properties. Proc. Natl. Acad. Sci. USA 97(11):5818-5823.

10. Park S-H, Zarrinpar A, & Lim WA (2003) Rewiring MAP kinase pathways using alternative scaffold assembly mechanisms. Science 299(5609):1061-1064.

11. Wang Y & Dohlman HG (2004) Pheromone signaling mechanisms in yeast: a prototypical sex machine. Science 306(5701):1508-1509.

12. Chapman SA & Asthagiri AR (2009) Quantitative effect of scaffold abundance on signal propagation. Mol. Syst. Biol. 5:313.

13. Yang J & Hlavacek WS (2011) Scaffold-mediated nucleation of protein signaling complexes: elementary principles. Math. Biosci. 232(2):164-173.

14. Suderman R & Deeds EJ (2013) Machines vs. Ensembles: Effective MAPK Signaling through Heterogeneous Sets of Protein Complexes. PLoS Comp. Biol. 9(10):e1003278.

15. Shao D, Zheng W, Qiu W, Ouyang Q, & Tang C (2006) Dynamic studies of scaffold-dependent mating pathway in yeast. Biophys. J. 91(11):3986-4001.

16. Thomson TM, et al. (2011) Scaffold number in yeast signaling system sets tradeoff between system output and dynamic range. Proc. Natl. Acad. Sci. USA 108(50):20265-20270.

17. Mayer BJ, Blinov ML, & Loew LM (2009) Molecular machines or pleiomorphic ensembles: signaling complexes revisited. J. Biol. 8(9):81.

18. Hu J, et al. (2011) Mutation that blocks ATP binding creates a pseudokinase stabilizing the scaffolding function of kinase suppressor of Ras, CRAF and BRAF. Proc. Natl. Acad. Sci. USA 108(15):6067-6072.

19. Hlavacek WS, et al. (2006) Rules for Modeling Signal-Transduction Systems. Sci. STKE 2006(344):re6-re6.

20. Danos V, et al. (2007) Rule-Based Modelling of Cellular Signalling. Lecture Notes in Computer Science, (Springer Berlin Heidelberg, Berlin, Heidelberg), Vol 4703, pp 17-41.

21. Danos V, Shao Z, Feret J, Fontana W, & Krivine J (2007) Scalable Simulation of Cellular Signaling Networks. Lecture Notes in Computer Science, (Springer Berlin Heidelberg, Berlin, Heidelberg), Vol 4807, pp 139-157.

22. Gillespie DT (1977) Exact stochastic simulation of coupled chemical reactions. J. Phys. Chem. 81.

23. Deeds EJ, Krivine J, Feret J, Danos V, & Fontana W (2012) Combinatorial complexity and compositional drift in protein interaction networks. PloS One 7(3):e32032.

24. Rowland MA, Fontana W, & Deeds EJ (2012) Crosstalk and competition in signaling networks. Biophys. J. 103(11):2389-2398.

25. McCaffrey G, Clay FJ, Kelsay K, & Sprague G (1987) Identification and regulation of a gene required for cell fusion during mating of the yeast Saccharomyces cerevisiae. Mol. Cell. Biol. 7(8):2680-2690.

26. Goldbeter A & Koshland DE (1981) An amplified sensitivity arising from covalent modification in biological systems. Proc. Natl. Acad. Sci. USA 78(11):6840-6844.

27. Suderman R, Bachman JA, Smith A, Sorger PK, & Deeds EJ (2017) Fundamental tradeoffs between information flow in single cells and cellular populations. Proc. Natl. Acad. Sci. USA 114(22):5755-5760.

28. Lin J, Harding A, Giurisato E, & Shaw AS (2009) KSR1 modulates the sensitivity of mitogen-activated protein kinase pathway activation in T cells without altering fundamental system outputs. Mol. Cell. Biol. 29(8):2082-2091.

29. Kirouac DC, et al. (2012) Creating and analyzing pathway and protein interaction compendia for modelling signal transduction networks. BMC Sys. Biol. 6(1):29.

30. Rowland MA, Greenbaum JM, & Deeds EJ (2017) Crosstalk and the evolvability of intracellular communication. Nat. Comm. 8:16009.

31. Bardwell L (2006) Mechanisms of MAPK signalling specificity. Biochem. Soc. Trans. 34(5):837.

32. Danos V, et al. (2012) Graphs, Rewriting and Pathway Reconstruction for Rule-Based Models. Leibniz International Proceedings in Informatics, eds D’Souza D, Kavitha T, & Radhakrishnan J (Leibniz International Proceedings in Informatics), pp 276-288.

33. Bardwell L (2005) A walk-through of the yeast mating pheromone response pathway. Peptides 26(2):339-350.

34. Bashan A & Yonath A (2008) Correlating ribosome function with high-resolution structures. Trends Microbiol. 16(7):326-335.

35. Swain PS, Elowitz MB, & Siggia ED (2002) Intrinsic and extrinsic contributions to stochasticity in gene expression. Proc. Natl. Acad. Sci. USA 99:12795-12800.

36. Sato PM, Yoganathan K, Jung JH, & Peisajovich SG (2014) The robustness of a signaling complex to domain rearrangements facilitates network evolution. PLoS Biol 12(12):e1002012.

37. Peisajovich SG, Garbarino JE, Wei P, & Lim WA (2010) Rapid Diversification of Cell Signaling Phenotypes by Modular Domain Recombination. Science 328(5976):368-372.

38. Gerhart J & Kirschner M (2007) The theory of facilitated variation. Proc. Natl. Acad. Sci. USA 104(suppl 1):8582-8589.

39. McCann JJ, Choi UB, & Bowen ME (2014) Reconstitution of multivalent PDZ domain binding to the scaffold protein PSD-95 reveals ternary-complex specificity of combinatorial inhibition. Structure 22(10):1458-1466.

